# The influence of intergenerational transfer of white matter tracts on early reading development

**DOI:** 10.1101/2020.10.09.333096

**Authors:** Maaike Vandermosten, Klara Schevenels, Maria Economou, Fumiko Hoeft

## Abstract

Parents have large genetic and environmental influences on their children’s cognition, behavior, and brain. Previous studies have indicated intergenerational transfer of the behavior of reading. Despite a close coupling between brain and behavior however, the intergenerational transfer of reading-related structural brain networks have not been investigated. Therefore, we investigated its parent-child associations for the first time. We examined how white matter tracts i.e., Arcuate Fasciculus (AF) and Inferior Fronto-Occipital Fasciculus (IFOF), are associated with children’s early reading development from Kindergarten to Grade 3 in 33 families. First, we observed in our sample of children -who all had typical reading skills despite half of them having an increased risk for dyslexia- that fractional anisotropy in bilateral IFOF and right AF correlated with reading development. Second, parent-child correlations were observed for bilateral IFOF but not for AF. Finally, we demonstrated that the relation between children’s IFOF and reading development was largely explained by parental IFOF. The findings preliminarily suggest that white matter organization in IFOF represents a pre-existing protective factor in children at risk, as it is mainly determined by biological parental factors. Large-scale intergenerational, multi-level and longitudinal studies are needed to understand the dynamic interrelations between brain, environment and behavior.

## Introduction

Intergenerational transfer of reading skills, measured by an association between reading skills of parents and their children, have been demonstrated in multiple studies (e.g. van Bergen et al. 2016; Vandermosten et al. 2016), yet the underlying mechanism of this parent-to-child transmission is not yet understood. According to the intergenerational multiple deficit model (van Bergen, van der Leij, de Jong, Bergen, & Leij, 2014), both parents pass on a liability for developmental dyslexia (i.e., a developmental disorder characterized by severe and persistent reading problems) by interwoven genetic and environmental pathways. This means that parents can contribute to their child’s reading skills via the environment they provide as well as via genetic and neurobiological contributions.

As for environmental factors, socio-economic status (SES) has been shown to be associated with children’s reading ability (e.g. Davis-Kean, 2005) as well as with white matter organization in the reading network (Ozernov-Palchik et al., 2019; Ursache & Noble, 2016; Vandermosten, Cuynen, Vanderauwera, Wouters, & Ghesquière, 2017). However, although SES is considered an environmental factor, there is a substantial genetic component to SES (Trzaskowski et al., 2014), and this might drive the link with reading. Indeed, twin-family studies on the intergenerational transfer of reading (Swagerman et al., 2017) suggest a potentially greater role for genetic or biological than for environmental factors. Based on large-scale diffusion MRI studies in twins and siblings, we know that white matter organization, quantified by fractional anisotropy (FA), is highly heritable when averaged across the whole-brain (heritability rate h^2^ > .75) (Chiang et al., 2011; Kochunov et al., 2015). Similar high heritability rates are observed for specific white matter tracts, such as the Arcuate Fasciculus (AF)/Superior Longitudinal Fasciculus (SLF) and the Inferior-Fronto-Occipital-Fasciculus (IFOF) (Kochunov et al., 2015), which have been found to be related to reading and reading-related cognitive skills (Gullick & Booth, 2014; Saygin et al., 2013; Vanderauwera, Wouters, Vandermosten, & Ghesquière, 2017; Vandermosten et al., 2012; Wang et al., 2017; Yeatman & Dougherty, 2012).

Yet, testing the heritability of neural structures associated with reading is not sufficient to implicate them as intergenerational neural mediators of reading. This would require intergenerational neuroimaging studies in which it is investigated whether similarities in neural systems of parents and their children mediate the relations to reading ability. To date, intergenerational brain studies have been conducted in the field of mental health, with a particular focus on corticolimbic circuitry, mood and anxiety (for a review see Ho, Sanders, Gotlib, & Hoeft, 2016). Regarding academic achievement, such as reading performance, studies have linked parental reading to the neurobiology of the child (Black et al., 2012; Farah, Dudley, Hutton, & Horowitz-Kraus, 2020; Vandermosten et al., 2017). A more direct link between parents’ and their children’s neural reading network however, remains to be investigated.

The current study therefore strives to investigate intergenerational associations of the neural reading network and to what extent this mediates reading development in children. With regard to the intergenerational correlation in the neural reading network, we select the dorsal pathway represented by the left AF and the ventral pathway represented by the left IFOF. These tracts have shown correlations with reading or reading-related cognitive skills across development, namely in pre-readers (Kraft et al., 2016; J. Vanderauwera et al., 2017; Wang et al., 2016), school-aged children (Gullick & Booth, 2014; Vanderauwera et al., 2018, 2017; Yeatman & Dougherty, 2012), and adults (Vandermosten et al., 2012). Yet, in contrast to adults, correlations in pre-readers and beginning readers are not only found in the left but also in the right AF and IFOF (Vandermosten, Hoeft, & Norton, 2016). More specifically, studies in pre-to beginning readers show that right hemispheric white matter tracts show decreased FA in children who later develop reading problems (Vanderauwera et al., 2017), as well as increased FA in the children that develop typical reading skills despite being at risk for dyslexia (Yu, Zuk, & Gaab, 2018). The latter suggests potential protective or compensatory right hemispheric systems in reading development, with at-risk children being resilient to reading problems by a greater involvement of the right hemisphere. Therefore, we investigate intergenerational correlations in left and right AF and IFOF in relation to early reading development (i.e., basic reading skills across the first years of reading instruction). More specifically, in a sample of 33 third grade children and their parents in the U.S., we conduct: (1) correlations between children’s FA in the selected white matter tracts and their reading development (Kindergarten to Grade 3); (2) intergenerational (i.e., parent-child) correlations in the selected white matter tracts; and (3) mediation analyses to examine to what extent the relation between children’s white matter tracts and reading development is mediated by parental white matter tracts.

## Results

### Reading development – white matter correlations in children

We first examined the association between children’s reading development and AF. In left AF, Bayesian statistics showed substantial evidence for no association between reading development and FA of left AF (*BF*_*10*_ = 0.31). On the other hand, there was strong evidence for an association between reading development and right AF (*BF*_*10*_ = 12.91). Frequentist analyses confirmed this by showing no significant correlations for left AF (*r* = .16, *p* = .39), but significant correlations for right AF (*r* = .50, *p* < .01).

Regarding the association between children’s reading development and IFOF, Bayesian statistics showed anecdotal evidence for an association between reading development and FA in left IFOF (*BF*_*10*_ = 1.1), and substantial evidence for an association in right IFOF (*BF*_*10*_ = 7.1). Frequentist analyses confirmed this pattern with correlations between reading development and left IFOF that did not reach significance (*r* = .32, *p* = .07) whereas correlations with right IFOF were significant (*r* = .46, *p* = .01).

In summary, Bayesian and frequentist analyses showed correlations (defined as *BF*_*10*_ > 3 and *p* < .05) for the relationship between children’s reading development and FA in right AF and right IFOF (see Figure 1), but only a trend for left IFOF and no evidence for a correlations for left AF.

**Figure 1:**
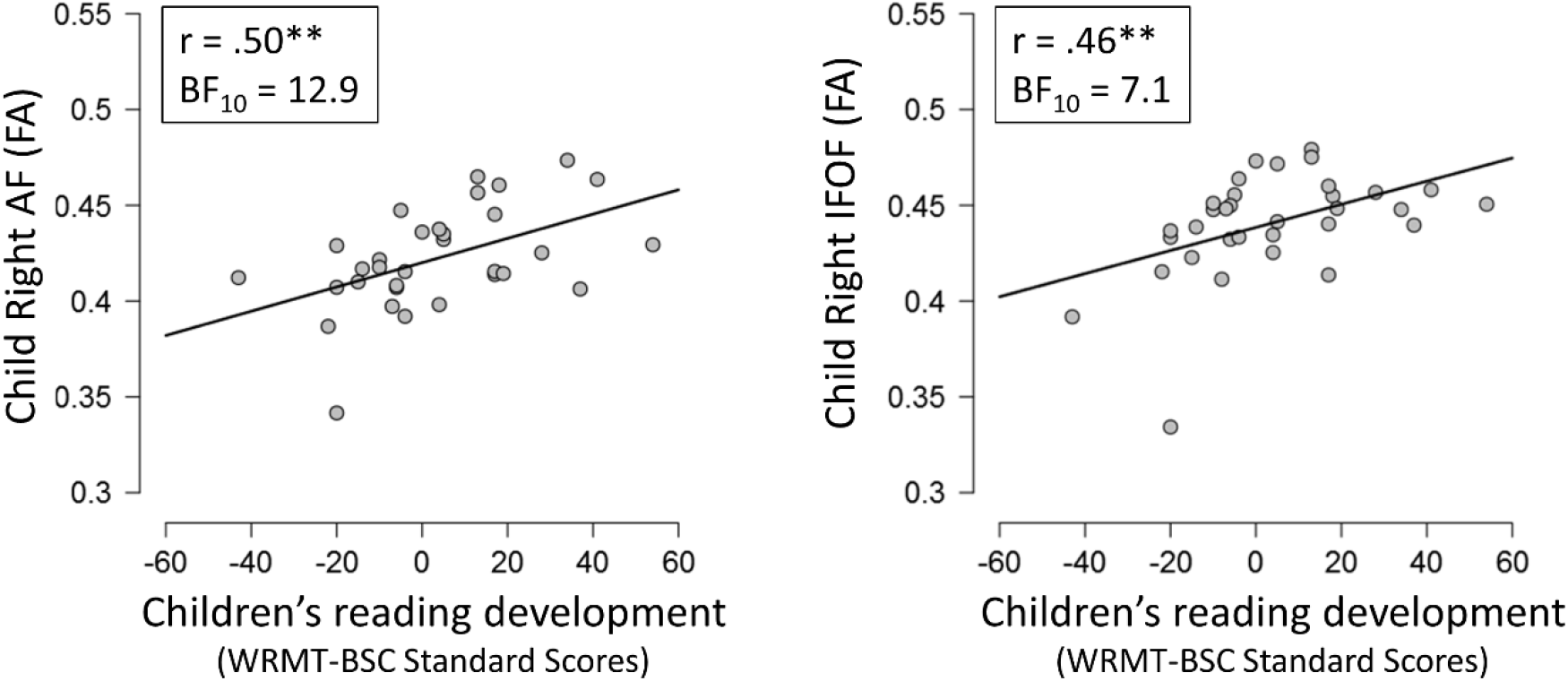
Correlations between children’s reading development and fractional anisotropy (FA) in right Arcuate Fasciculus (AF) and Inferior-Fronto-Occipital-Fasciculus (IFOF). Reading development was defined as change in standard scores (SS) of Basic Skill Cluster of the Woodcock Reading Mastery Tests (WRMT-BSC) from Time point 1 in kindergarten to Time point 2 in third grade. Since we report in SS, a change score of 0 indicates they showed typical development. A positive score indicates change that was greater than what is typically expected. For results using raw scores, please see Supplementary Information. Only correlations for which Bayesian statistics showed at least substantial evidence (BF > 3) and frequentist statistics showed significant results (p < .05) are depicted. * p < .05; ** p < .01, *** p < .001

### Intergenerational correlations of FA in white matter tracts

For each white matter tract, i.e., left and right AF and IFOF, Pearson correlations were calculated between parents’ and their child’s FA using both Bayesian and frequentist approaches. In general, father and child, and mother and child correlations were observed for FA in bilateral IFOF but not for either left or right AF.

More specifically, for left AF, Bayesian factors were all below 1, indicating to be in favor of no association (child – mother: *BF*_*10*_ = .26; child – father: *BF*_*10*_ = .43). Frequentist analyses showed no significant correlations (child – mother: *r* = −.01, *p* = .97; child – father: *r* = .23, *p* = .32). For right AF, Bayesian factors were again below 1 (child – mother: BF_10_ = .28; child – father: BF_10_ = .55). Frequentist analyses also showed no significant correlations (child – mother: *r* = −.07, *p* = .76; child – father: *r* = −.28, *p* = .23).

For left IFOF, Bayesian analyses indicated that there was anecdotal evidence for an association between children’s and mothers’ FA (*BF*_*10*_ = 1.7), and very strong evidence for an association between children’s and fathers’ FA (*BF*_*10*_ = 47.0). Using frequentist analyses, both correlations, children’s and mothers’ FA (*r* = .42, *p* = .04) and children’s and fathers’ FA (*r* = .67, *p* < .001) reached significance. For right IFOF, Bayesian statistics revealed that there was substantial evidence (*BF*_*10*_ = 6.5) for an association between children’s and mothers’ FA, and that there was anecdotal evidence for an association between children’s and fathers’ FA (*BF*_*10*_ = 1.3). Frequentist analyses showed a significant correlation between children’s and mothers’ FA (*r* = .53, *p* = .01), and children’s and fathers’ FA showed a trend for significant correlation (*r* = .41, *p* = .07). The intergenerational correlations for IFOF are depicted in Figure 2.

**Figure 2:**
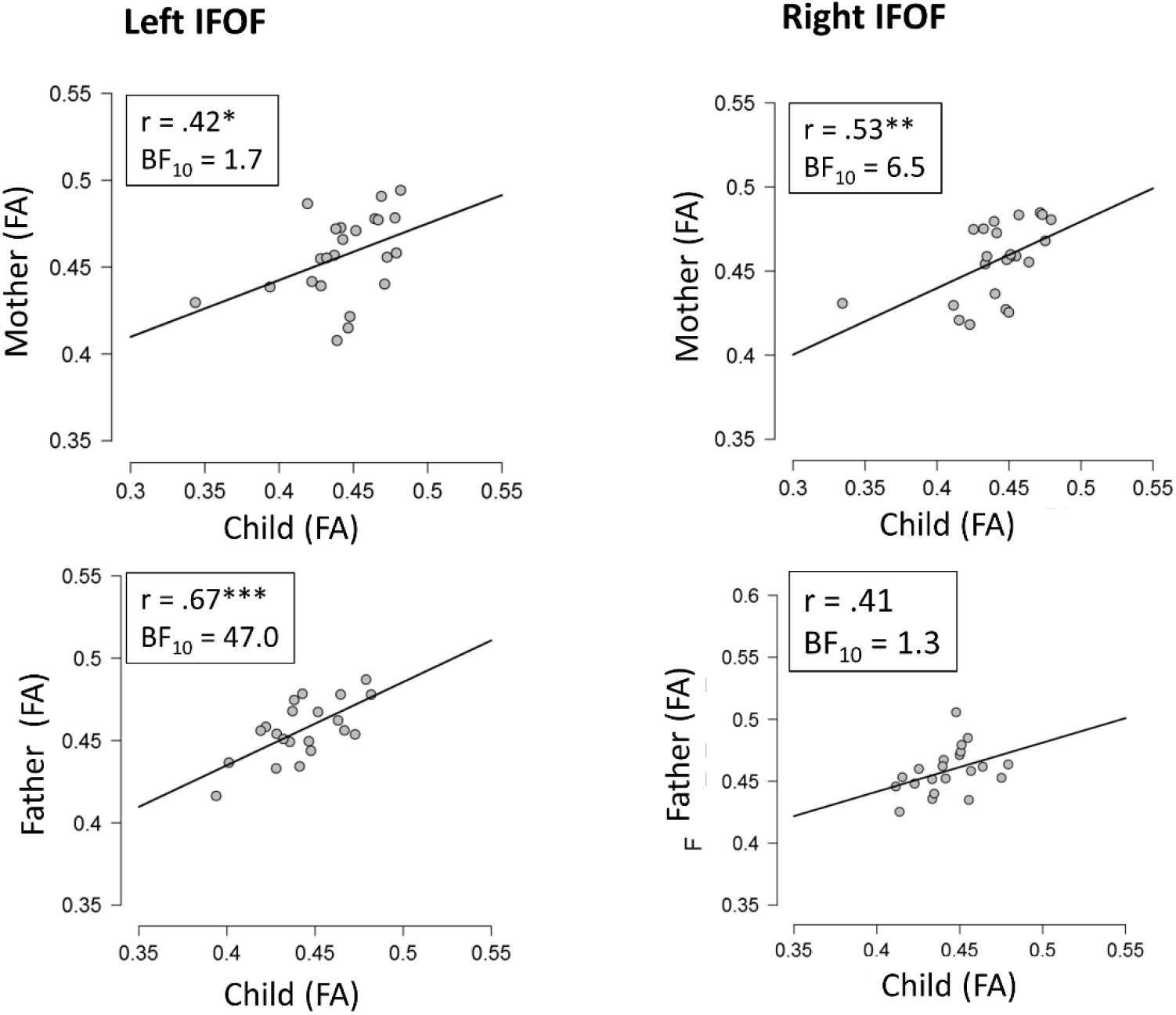
Intergenerational transfer of fractional anisotropy (FA) in ventral Inferior Fronto-Occipital-Fasciculus (IFOF). * p < .05; ** p < .01, *** p < .001

### Mediation Analyses

Given that intergenerational correlations were found for left and right IFOF, we conducted mediation analyses with parental IFOF as a potential mediator for the relation between a child’s IFOF and reading development. More specifically, children’s reading development was selected as the dependent variable, children’s FA in IFOF as the independent variable, and mothers’ and fathers’ FA in IFOF as mediators (see Figure 3). Analyses were run separately for left and right IFOF.

**Figure 3:**
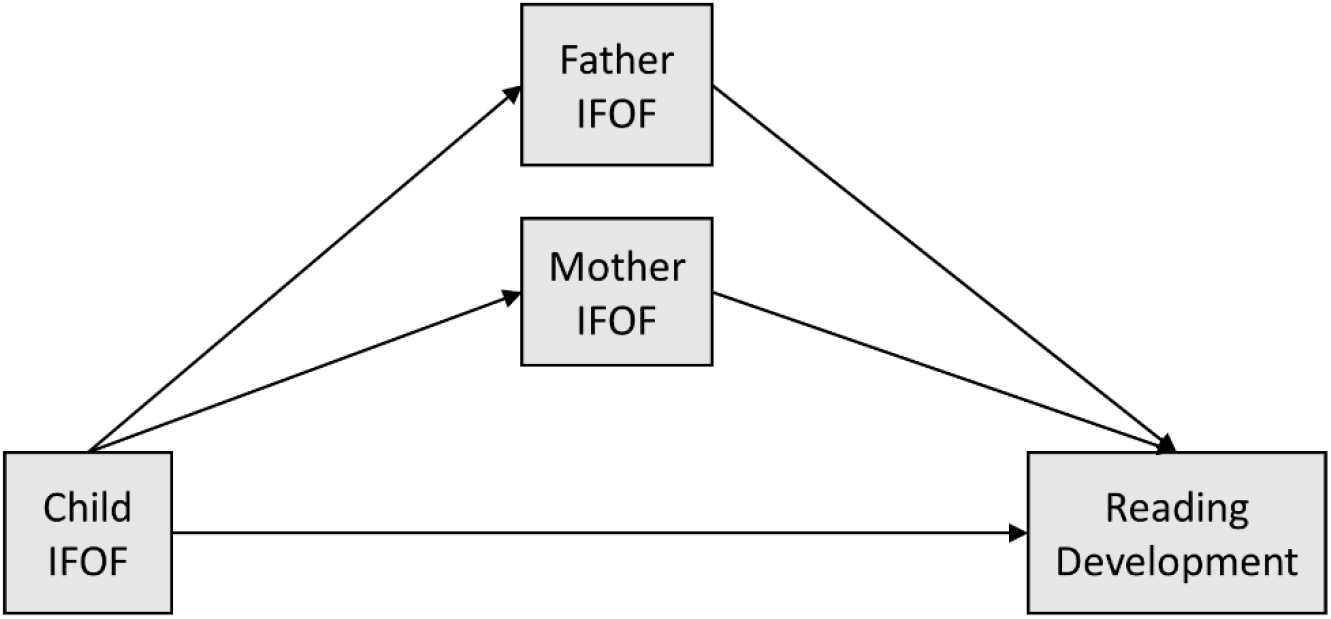
A schematic diagram of mediation effects of parental IFOF on the relation between children’s IFOF and reading development.

With regards to the right IFOF, the total effect, i.e., the regression coefficient between children’s right IFOF FA and reading development, was 16.89 and significant (*z* = 2.98, *p* < .01), which is in line with the correlation analyses reported above. However, when the mediation effect (i.e., parental FA in right IFOF) was taken into account, the direct effect of child’s right IFOF on reading development decreased to 3.14 and became non-significant (*z* = .24, *p* = .81). The indirect effect was (18)*(0.38) = 6.81 for mothers and (25)*(0.28) = 6.94 for fathers. These results imply that the indirect effects together (13.75) explained 81% of the total effect of children’s FA in right IFOF on reading development. However, when we tested the significance of these indirect effects using bootstrapping procedures (unstandardized indirect effects were computed for each 1,000 bootstrapped sample), the indirect effect was non-significant (*p* = .25; 95% confidence interval ranged from −12.84 to 49.06).

Regarding the left IFOF, the total effect, i.e., the regression coefficient between children’s left IFOF and reading development, was 11.28 and close to significant (*z* = 1.96, *p* = .05], which is in line with the trend reported for the correlation analyses above. However, when the mediation effect (i.e., parental FA in left IFOF) was taken into account, the direct effect of children’s left IFOF on reading development decreased to 2.74 and became non-significant (*z* = .24, *p* = .81). The indirect effect was (14)*(0.15) = for mothers and (29)*(0.23) = 6.6 for fathers. These results imply that the indirect effects together (8.54) explained 75% of the total effect of child’s left IFOF on reading development. However, when we tested the significance of these indirect effects using bootstrapping procedures (unstandardized indirect effects were computed for each 1,000 bootstrapped sample), the indirect effect was non-significant (*p* = .39; 95% confidence interval ranged from −10.95 to 28.03).

## Discussion

In this study, we aimed to investigate whether associations in white matter organization of reading-related tracts between parents and their children can explain the amount of reading progress a child makes during the first years of formal schooling. In our sample of children that consisted of typical reading children in the U.S. with and without a family risk for dyslexia, white matter organization of right IFOF and right AF, and to some extent left IFOF, correlated with how much reading developed over 3 years. In addition, we observed intergenerational transfer of white matter organization in bilateral IFOF, but no evidence was found for intergenerational transfer of bilateral AF. Finally, mediation analyses showed that the association between children’s IFOF and reading development was largely (71% to 81%) explained by parental IFOF, suggesting a mediating role by parents’ neurobiology, although these indirect effects did not reach significance.

Learning to read requires practice and instruction because a child needs to construct a neural network dedicated to reading by connecting and reorganizing the neural systems for language and visual perception (Dehaene et al., 2010). White matter tracts are key to build up such a network as they consist of bundles of myelinated axons allowing communication between distant cortical regions. In a matured reading network of adults, left hemispheric tracts are dominant for reading, with left AF involved in the phonological aspects, and left IFOF in the orthographic aspects of reading (Vandermosten et al., 2012). However, in children at the onset of reading acquisition both left and right structures seem to play a key role in reading (Vandermosten et al., 2016). At a pre-reading stage, bilateral dorsal and ventral tracts are important for reading-related cognitive skills, and neural specialization and left dominance only emerge after years of reading instruction (Vanderauwera et al., 2018). Both left and right tracts are also important for the early prediction of dyslexia. Several independent studies have shown that structural organization in left AF predicts which pre-readers will subsequently develop reading problems (Kraft et al., 2016; Langer et al., 2015; Vanderauwera et al., 2017; Wang et al., 2017). Yet, white matter organization of AF in the right hemisphere, especially the anterior segment connecting frontal and parietal regions, appear to play an important role in protecting high-risk children from developing dyslexia (Wang et al., 2017; Yu et al., 2018; Zuk et al., 2020). The development of well-organized right hemispheric tracts in order to overcome the risk for dyslexia could explain the correlational results we obtained; namely the better the right AF and IFOF were organized at Grade 3, the more reading improvement occurred during the first years of reading acquisition. The observation that our sample included at-risk children who eventually all developed reading skills within the typical range suggests that well-structured right hemispheric tracts seem to have protected them from developing reading problems. We also observed a weak association between reading development and left IFOF, which is not in line with previous studies showing mainly protective mechanisms in the right hemisphere. Yet, given the weak association, further replication is needed before conclusions can be drawn on whether left ventral tracts could also play a protective role.

In a second step, in order to understand the underlying mechanisms of the association between a child’s right hemispheric tracts and reading development, we investigated whether parent-child correlations of white matter organization could be observed, and whether parental white matter tracts could explain the relationship between children’s white matter tracts and their reading development. According to the intergenerational multiple deficit model (van Bergen et al., 2014), both parents influence a child’s behavioral via intertwined genetic and environmental pathways. These pathways pass through neural intermediaries of the parents and the child, yet direct parent-child associations at the neural level have not been investigated. In our sample consisting of parent-child data of 33 families, we observed intergenerational correlations for IFOF, which was expected based on the high heritability rates of FA in white matter tracts, including IFOF, that were observed in twin/sibling studies (Peter Kochunov et al., 2015). For left IFOF, we found strong correlations between fathers and children, and for right IFOF, strong correlations between mothers and children, although similar trends were also observed for the other parent. Hence, although sex-specific intergenerational effects have been reported for corticolimbic system in relation to mood regulation (Yamagata et al., 2016), our current data suggest no paternal or maternal specific transfer for reading-related tracts. As an important next step, we examined if FA of parental IFOF mediates the relationship between children’s IFOF and reading development. Our results demonstrate a mediating role of parental IFOF since the relation between a children’s IFOF and reading development was for the vast majority (75% - 81%) explained by parental IFOF. We must however, interpret our findings cautiously given that the indirect effects did not reach significance in our rather small sample. Nevertheless, since a large majority of the total effect was explained by the indirect effects (i.e., mediators), our results tentatively suggest that the organization of a child’s right IFOF might prevent a child at risk for dyslexia to develop reading problems, and that this protective mechanism is transferred via the parents, hence might be in place from birth. Hence, this suggests that a reliance on right hemispheric tracts does not emerge in response to reading difficulties, i.e., as a compensatory mechanism, but rather before the onset of reading problems, i.e., as a protective mechanism.

Finally, a different pattern of correlations for AF and IFOF was observed regarding intergenerational transfer. While evidence was found for intergenerational correlations in bilateral IFOF, no intergenerational correlations were found for bilateral AF. The lack of intergenerational correlations in AF was surprising given diffusion imaging data from large scale twin and sibling studies demonstrating high heritability rates for FA in both AF and IFOF (.70 – .90) (Kochunov, Jahanshad, Marcus, & Winkler, 2015). However, these studies utilize twin and sibling data who share their family environment and this might results in inflated heritability rates (Blangero et al., 2013). One potential explanation for the observed intergenerational correlations in our study is that family environment could impact AF more than IFOF, hence resulting in less clear intergenerational transfer of AF organization due to environmental influences, and a more direct transfer of parents’ neurobiology for IFOF. This explanation may be in line with developmental neuroimaging studies showing that IFOF is formed and organized already early life (Brauer, Anwander, Perani, & Friederici, 2013), while during the first years of schooling FA development is more pronounced in AF than in IFOF (Dimond et al., 2020), hence allowing for environmentally-induced plasticity in the AF during child development. Previous studies on SES, which is the most commonly used environmental factor, indicate mixed findings, with studies showing SES impact on AF (Ozernov-Palchik et al., 2019), on IFOF (Gullick, Demir-Lira, & Booth, 2016; Vandermosten et al., 2017), or on both (Dufford & Kim, 2017). However, SES is not a pure environmental factor and also contains large genetic contributions (Trzaskowski et al., 2014). Moreover, environmental factors that are more directly related to literacy development might be more informative. From that perspective, recent studies on language (Romeo et al., 2018) and literacy (Su et al., 2020) exposure during the first years of life show a substantial impact of language/literacy exposure on dorsal and less on ventral tracts. Moreover, the impact of early literacy exposure on AF is long lasting and still present in the more matured brain of adolescence (Su et al., 2020). Although in our study, we have no direct measures of early language and literacy exposure to confirm this hypothesis, future studies should further investigate this by including data from online recordings of early language and literacy exposure or by investigating the effects of early reading and language interventions on white matter tracts in addition to intergenerational diffusion MRI data. A second potential explanation of why no intergenerational correlations were found for AF while we did for IFOF is that parents each provide genetic contributions to the development of their children’s white matter tract and that the resemblance between parents’ and children’s tract might be larger when fathers’ and mothers’ tracts are alike. Although assortative mating, i.e., tendency to choose partners that are similar to themselves in phenotypic characteristics, has been demonstrated for reading skills, it has not been investigated at the endophenotypic level, i.e., similarities between partners in white matter tracts. To examine this potential explanation, we have post-hoc calculated correlations between white matter tracts of mothers and fathers and we found assortative mating for left IFOF (BF_10_ = 3.134, r = .543, p = .024), but no significant correlations for bilateral AF nor right IFOF (BF_10_ < .34, p > .610). Hence, a higher resemblance among parents cannot fully explain why more parent-child similarities are found for IFOF and not for AF, since right IFOF showed no assortative mating but did show intergenerational correlations. A final explanation for the lack of intergenerational correlations for AF is the relatively small sample size which might have led to insufficient power to observe intergenerational correlations. Traditional p value null-hypothesis significance testing (i.e., frequentist analyses) is arguably appropriate to quantify evidence against the null-hypothesis (i.e., presence of an effect) but cannot interpret non-significant findings in terms of whether there is evidence for absence of an effect or whether results are simply not informative due to insufficient power to detect an effect (Keysers, Gazzola, & Wagenmakers, 2020). Therefore, we have additionally applied Bayesian hypothesis testing which can help in interpreting non-significant results, even in small samples (Keysers et al., 2020). Our results show that for AF the Bayes factor of mother-child intergenerational correlations were below 1/3 (left BF_10_ = .26; right BF_10_ = .28), which allowed us to conclude that there is substantial evidence for the absence of mother-child correlations for AF rather than being due to insufficient power to detect a correlation. However, the Bayes factor of father-child intergenerational correlation for AF are between 1/3 and 1 (left BF_10_ = .43; right BF_10_ = .55), which indicates that there is insufficient evidence to draw conclusion for or against the null hypothesis (Keysers et al., 2020). Hence, at present, although we can conclude that our results indicate that there is no mother-child association in AF, large-scale studies are needed to confirm the lack of correlations for father-child in AF.

To conclude, our results indicate that white matter organization of right hemispheric reading tracts play an important role in how much reading progress a child makes during the first years of reading acquisition, especially in children at risk for dyslexia. Right IFOF seems to be involved in protective mechanisms that is transferred from parent to child, given that intergenerational correlations are found for bilateral IFOF. Moreover, parental IFOF explains the vast majority of the variance of the relationship between children’s IFOF and reading development, indicating the importance of parental neurobiology in understanding the underlying mechanisms. The lack of intergenerational correlations for AF requires further investigation, but one hypothesis is that the dorsal tract, which is in contrast to ventral tracts, is still immature during the first years of life, and is more sensitive to plasticity induced by the amount and quality of language and literacy exposure. Intergenerational and longitudinal neuroimaging studies that include large samples and use ecologically valid early environmental measures are needed to further investigate the complex intertwined relationships between reading, brain and environment.

## Methods

### 1. Participants

Diffusion MRI data were collected in 35 children at Grade 3, which is a subset of an original 51 participants who took part in a larger study (Black et al., 2012; Gimenez et al., 2014; Hosseini et al., 2013; for details see Myers et al., 2014). Usable diffusion MRI and T1 data were available in 33 children (for details see Table 1a), 23 mothers and 21 fathers (MRI data of both parents were available in 17 children, of only mothers in 6 children, of only fathers in 4 children and of no parents in 6 children) (for details see Table 1b). Of the 33 children included, 19 had a family risk for dyslexia, defined as having one first degree relative with dyslexia using the Adult Reading History Questionnaire (ARHQ) with a score of .4 or more in at least one biological parent (Lefly & Pennington, 2000). This oversampling of family risk children relative to the general population increased the chance of including children that subsequently developed dyslexia since the prevalence rate for dyslexia in family risk children is 40-60% while it is only 5-10% in a typical sample (Gilger, Pennington, & Defries, 1991). However, despite the elevated risk in more than half of the participants, reading tests (i.e., Basic Skill Cluster [BSC] score of Woodcock Reading Mastery Test - Revised Normative Update [WRMT-R/NU], which is a combination of Word Identification [WID] and Word Attack [WA] subtests at Grade 3, showed that all children performed either average or above average (for *M* and *SD* see Table 1a). This implies that our sample is atypical in its prevalence for reading problems, suggesting that it contains readers who are resilient to their risk for dyslexia or have compensated for their reading problems. Children had no neurological or psychiatric disorders including attention deficit hyperactivity disorder (ADHD), were not on any medication, and had no contraindications to MRI, as was reported by the parents. The study was approved by the Stanford University Panel on Human Subjects in Medical Research and University of California, San Francisco Human Research Protection Program. Informed consent for all children was obtained from a parent (verbal assent was given by the child) and informed consent for all parents was provided by themselves.

**Table 1a:** Data on demographic, diffusion MRI and behavioral testing in children

|  | Child |
| --- | --- |
| Handedness (R/L) | 27/6 |
| Family Risk (low/high) | 14/19 |
| Gender (F/M) | 12/21 |
| **Diffusion MRI (Grade 3)** |  |
| N | 33 |
| Age at MRI-scan | 8.27 (.48) |
| L AF (FA) | .428 (.021) |
| R AF (FA) | .423 (.027) |
| L IFOF (FA) | .441 (.029) |
| R IFOF (FA) | .440 (.027) |
| **Reading assessment (Grade 3)** |  |
| N | 33 |
| Age at testing | 8.21 (.46) |
| WRMT-BSC<sup>1</sup> | 118.8 (16.5) |
| - WRMT-WID<sup>1</sup> | 115.9 (12.6) |
| - WRMT-WA<sup>1</sup> | 113.7 (15.2) |
| **Reading assessment (Kindergarten)** |  |
| Age at testing | 5.56 (0.46) |
| WRMT-BSC<sup>1</sup> | 115.4 (27.0) |
| - WRMT-WID<sup>1</sup> | 120.1 (32.2) |
| - WRMT-WA<sup>1</sup> | 111.0 (14.4) |
| **Reading development (Grade 3 - Kindergarten)** |  |
| WRMT_BSC<sup>1</sup> | 3.39 (20.84) |
| - WRMT_WID<sup>2</sup> | 51.8 (13.4) |
| - WRMT_WA1<sup>2</sup> | 20.8 (8.2) |

<sup>1</sup> Standardized reading scores with $M = 100$ and $SD = 15$ ; <sup>2</sup> Raw scores; IFOF = Inferior Fronto-Occipital Fasciculus; AF = Arcuate Fasciculus

**Table 1b:** Data on demographic, diffusion MRI and behavioral testing in parents

|  | Mother | Father |
| --- | --- | --- |
| **Diffusion MRI testing** |  |  |
| N | 23 | 21 |
| Age at MRI-scan | 42.3 (4.2) | 45.49 (4.92) |
| L AF (FA) | .450 (.019) | .443 (.017) |
| R AF (FA) | .443 (.020) | .442 (.020) |
| L IFOF (FA) | .457 (.024) | .457 (.018) |
| R IFOF (FA) | .456 (.022) | .459 (.019) |

IFOF = Inferior Fronto-Occipital Fasciculus; AF = Arcuate Fasciculus

### 2. Reading assessment

In children, behavioral testing included a large test battery on, among others, reading (Test of Word Reading Efficiency, WRMT-R/NU), phonology (Comprehensive Test of Phonological Processing, Rapid Automatized Naming), language (Clinical Evaluation of Language Fundamentals 4 & Peabody Picture Vocabulary Test 4) and academic achievement (Woodcock-Johnson III Tests of Achievement), collected at different time points across reading development. In the current study, we focused on reading development between Kindergarten and Grade 3. Reading ability was defined by the Basic Skill Cluster of the Woodcock Reading Mastery Tests (WRMT-BSC), which is a composite score of the standard scores of the subtests Word Identification (WRMT-WID) and Word Attack (WRMT-WA). In order to investigate the individual reading development, the standardized scores of Basic Skill Cluster (WRMT-BSC) in Kindergarten were subtracted from the ones in Grade 3. Note that a reading development score of zero on the standardized scores of WRMT-BSC indicates that reading developed at an expected rate (i.e., staying at the same standardized score), while a positive or negative developmental score indicates a greater or less gains in reading relative to the general population, respectively. Analyses on reading development based on the raw data of the individual subtests (WRMT-WID and WRMT-WA) are provided in Supplementary Information and show the same pattern of results as when reading development is based on the standardized scores of WRMT-BSC. Details on the reading scores of the children at each time point and the reading development score can be found in Table 1a.

### 3. MRI data acquisition and tractography

Diffusion MRI scanning was done on a GE Healthcare 3.0 Tesla at the Richard M. Lucas Center for Imaging at Stanford University. To prepare children for the MRI scan, families received a CD of scanner noises and a DVD of a child going into a scanner, and participated in simulated MRI sessions at the center to further minimize movement. Images acquired included a high-angular resolution diffusion-imaging scans\ using the following parameters: 150 diffusion-encoding gradient directions, diffusion sensitivity of b = 2500 s/mm2, repetition time (TR) = 5000ms, echo time (TE) = 82.6 ms, matrix size 128 × 128, spatial in-plane resolution 2.0 × 2.0mm and slice thickness 3.0 mm. All pre-processing of the dMRI data was done using the ‘Explore DTI’ software (Leemans et al., 2009), consisting of visual quality assurance, and rigorous motion, eddy current-induced distortion and Eche Planar Imaging distortion correction. No participants were excluded due to excessive motion in scans, but for 2 of the 35 children T1 were not usable to perform EPI correction during preprocessing, hence resulting in a final dataset of 33 children. A diffusion tensor model was estimated using non-linear least square fitting. No normalization to a standard atlas took place. Whole-brain tractography was performed with the following parameters: FA-threshold of .20, maximum turning angle of 40 degrees, step length between calculations of 1 mm and fiber length range of 50-500 mm. Tractography of the white matter tracts of interest, i.e., bilateral AF and IFOF, was performed with the TrackVis software (Wang et al., 2007). All tracts were manually delineated for each subject using a region of interest (ROI) approach, based on anatomical landmarks in color-coded maps (Catani and Thiebaut de Schotten, 2008; Thiebaut de Schotten et al., 2011; Wakana et al., 2007). Given the different naming and definition for the AF (for more details see Dick & Trembley, 2012), we clarify more explicitly how we defined the tract. We used the ROIs as described in Wakana et al. (2007) for ‘SLF’, but used the terminology ‘AF’, in line with Catani et al (2008), since SLF is referring to tracts running more dorsally and difficult to delineate with the tensor model (see Zhang & Ramus, 2015). This implies that the AF, as we defined it, entails projections from frontal to parietal as well as to temporal areas. In Figure 4, an example of the AF and IFOF in one representative child is depicted. Per individual, we extracted the average FA of each of these 4 tracts. For all children and parents for whom dMRI data were available all four tracts could be delineated, except for one child for whom right AF could not be delineated.

**Figure 4:**
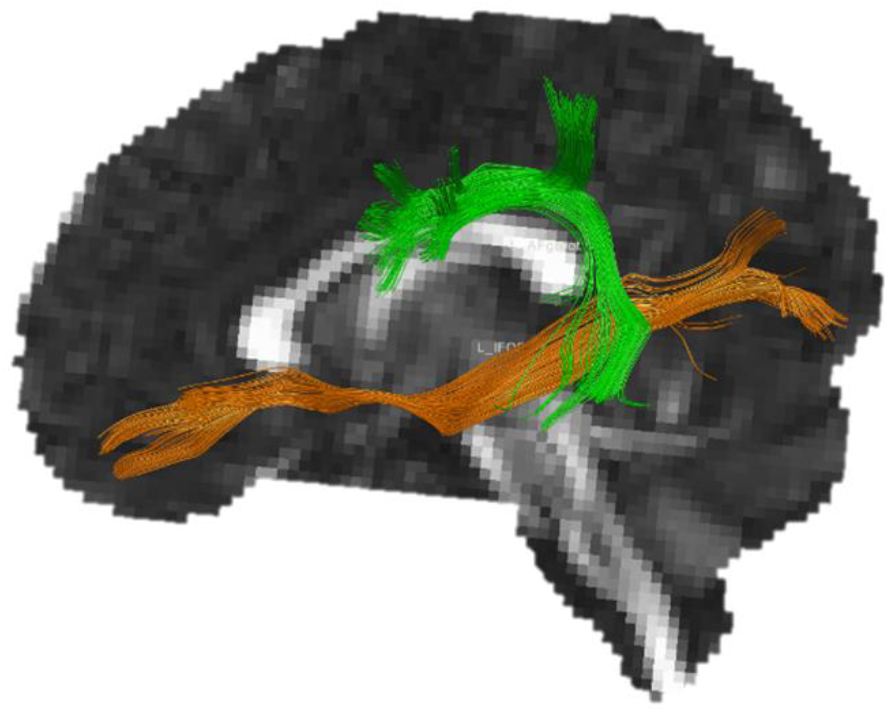
Example of left Arcuate Fasciculus (AF) in green and Inferior-Fronto-Occipital Fasciculus (IFOF) in orange for one child participant

### 4. Statistical analyses

Statistical analyses included correlational and mediation analyses, and were performed with the statistical software JASP (JASP Team, 2020). Concerning correlations, we calculated: (1) correlations between FA of the tracts of children and their reading development (between kindergarten and Grade 3), and (2) intergenerational white matter correlations, i.e., correlations between FA of the children’s tract and FA of the corresponding tract in their parents. We performed correlations using botha Bayesian and a frequentist approach. The advantage of Bayesian statistics is that one is able to quantify the amount of evidence that data provide for the null hypothesis as well as against it (Keysers et al., 2020). Correlations with Bayes factors (BF10) of 1, 1-3, 3-10, 10-30, 30-100, or > 100 respectively point towards no, anecdotal, substantial, strong, very strong, or decisive evidence for the hypothesis of an association between two variables (Jeffreys, 1961), whereas Bayes factors below 1 indicate that the null hypothesis of no association is more likely than the existence of an association. Results of frequentist approaches to statistical testing are also reported. For mediation analyses, we first selected the white matter tracts that showed intergenerational correlations, defined as correlations that provided at least substantial evidence using Bayesian statistics (BF_10_ > 3) or a significant correlation using frequentist statistics (p < .05). Then, via mediation analyses, we examined whether the relation between children’s reading development and children’s FA in the selected tract was mediated by mothers’ and fathers’ FA in the corresponding tract.

## Supporting information

For results using raw scores, please see Supplementary Information

## Acknowledgements

This study was supported by the Eunice Kennedy Shriver National Institute of Child Health and Human Development (NICHD) K23HD054720 (PI: F. Hoeft), Child Health Research Program (aka Lucile Packard Foundation for Children’s Health, Spectrum Child Health & Clinical and Translational Science Award) (PI: F. Hoeft). F. Hoeft is additionally supported by NIH R01HD078351 (PI: F. Hoeft), R01HD086168 (PIs: F. Hoeft & K. Pugh), P50HD52120 (PI: R. Wagner), R01HD044073 (PI: Cutting), R01HD096261 (PI: Hoeft), R01HD094834 (PIs: Hoeft & Hancock), NSF BCS-2029373 (PI: F. Hoeft), UCSF Dyslexia Center, Oak Foundation ORIO-16-012 & OCAY-19-215 (PI: F. Hoeft), and Lori and Ray DePole. The research was also supported by the postdoctoral and travel grant of Maaike Vandermosten (Research Foundation Flanders).

## Author contributions

F.H. designed the study and collected data with her students. F.H. and M.V. conceived the particular idea of the manuscript. Under supervision of M.V., K.S and M.E. did the manual tractography analyses. M.V. did the statistical analyses and took the lead in writing the manuscript. F.H., M.V, and K.S. provided critical feedback and helped shape the manuscript.

## Statement

We confirm that all methods were carried out in accordance with relevant guidelines and regulations.

## Notes

### Competing Interest Statement

The authors have declared no competing interest.

## References

1. Black, J. M., Tanaka, H., Stanley, L., Nagamine, M., Zakerani, N., Thurston, A., … Hoeft, F. (2012). Maternal history of reading difficulty is associated with reduced language-related gray matter in beginning readers. NeuroImage, 59(3), 3021–3032. https://doi.org/10.1016/j.neuroimage.2011.10.024

2. Blangero, J., Diego, V. P., Dyer, T. D., Almeida, M., Peralta, J., Kent, J. W., … GÖring, H. H. H. (2013). A Kernel of Truth. Statistical Advances in Polygenic Variance Component Models for Complex Human Pedigrees. In Advances in Genetics (Vol. 81, pp. 1–31). Academic Press Inc. https://doi.org/10.1016/B978-0-12-407677-8.00001-4

3. Brauer, J., Anwander, A., Perani, D., & Friederici, A. D. (2013). Dorsal and ventral pathways in language development. Brain and Language, 127(2), 289–295. https://doi.org/10.1016/j.bandl.2013.03.001

4. Chiang, M. C., McMahon, K. L., de Zubicaray, G. I., Martin, N. G., Hickie, I., Toga, A. W., … Thompson, P. M. (2011). Genetics of white matter development: A DTI study of 705 twins and their siblings aged 12 to 29. NeuroImage, 54(3), 2308–2317. https://doi.org/10.1016/j.neuroimage.2010.10.015

5. Davis-Kean, P. E. (2005). The influence of parent education and family income on child achievement: The indirect role of parental expectations and the home environment. Journal of Family Psychology, 19(2), 294–304. https://doi.org/10.1037/0893-3200.19.2.294

6. Dehaene, S., Pegado, F., Braga, L. W., Ventura, P., Nunes Filho, G., Jobert, A., … Cohen, L. (2010). How learning to read changes the cortical networks for vision and language. Science, 330(6009), 1359–1364. https://doi.org/10.1126/science.1194140

7. Dimond, D., Rohr, C. S., Smith, R. E., Dhollander, T., Cho, I., Lebel, C., … Bray, S. (2020). Early childhood development of white matter fiber density and morphology. NeuroImage, 210. https://doi.org/10.1016/j.neuroimage.2020.116552

8. Dufford, A. J., & Kim, P. (2017). Family income, cumulative risk exposure, and white matter structure in middle childhood. Frontiers in Human Neuroscience, 11, 547. https://doi.org/10.3389/fnhum.2017.00547

9. Farah, R., Dudley, J., Hutton, J., & Horowitz-Kraus, T. (2020). Maternal reading and fluency abilities are associated with diffusion properties of ventral and dorsal white matter tracts in their preschool-age children. Brain and Cognition, 140, 105532. https://doi.org/10.1016/j.bandc.2020.105532

10. Gilger, J. W., Pennington, B. F., & Defries, J. C. (1991). Risk for Reading Disability as a Function of Parental History in Three Family Studies (pp. 17–29). Springer, Dordrecht. https://doi.org/10.1007/978-94-011-2450-8_2

11. Gimenez, P., Bugescu, N., Black, J. M., Hancock, R., Pugh, K., Nagamine, M., … Hoeft, F. (2014). Neuroimaging correlates of handwriting quality as children learn to read and write. Frontiers in Human Neuroscience, 8, 155. https://doi.org/10.3389/fnhum.2014.00155

12. Gullick, M. M., & Booth, J. R. (2014). Individual differences in crossmodal brain activity predict arcuate fasciculus connectivity in developing readers. Journal of Cognitive Neuroscience, 26(7), 1331–1346. https://doi.org/10.1162/jocn_a_00581

13. Gullick, M. M., Demir-Lira, Ö. E., & Booth, J. R. (2016). Reading skill-fractional anisotropy relationships in visuospatial tracts diverge depending on socioeconomic status. Developmental Science, 19(4), 673–685. https://doi.org/10.1111/desc.12428

14. Ho, T. C., Sanders, S. J., Gotlib, I. H., & Hoeft, F. (2016). Intergenerational Neuroimaging of Human Brain Circuitry. https://doi.org/10.1016/j.tins.2016.08.008

15. Hosseini, S. M. H., Black, J. M., Soriano, T., Bugescu, N., Martinez, R., Raman, M. M., … Hoeft, F. (2013). Topological properties of large-scale structural brain networks in children with familial risk for reading difficulties. NeuroImage, 71, 260–274. https://doi.org/10.1016/j.neuroimage.2013.01.013

16. JASP Team,; (2020). No Title.

17. Jeffreys, H. (1961). Theory of probability (3rd ed.). Oxford: Oxford University Press.

18. Keysers, C., Gazzola, V., & Wagenmakers, E. J. (2020, July 1). Using Bayes factor hypothesis testing in neuroscience to establish evidence of absence. Nature Neuroscience. Nature Research. https://doi.org/10.1038/s41593-020-0660-4

19. Kochunov, P, Jahanshad, N., Marcus, D., & Winkler, A. (2015). Heritability of fractional anisotropy in human white matter: a comparison of Human Connectome Project and ENIGMA-DTI data. Neuroimage. Retrieved from http://www.sciencedirect.com/science/article/pii/S1053811915001512

20. Kochunov, Peter, Jahanshad, N., Marcus, D., Winkler, A., Sprooten, E., Nichols, T. E., … Van Essen, D. C. (2015). Heritability of fractional anisotropy in human white matter: A comparison of Human Connectome Project and ENIGMA-DTI data. NeuroImage, 111, 300–311. https://doi.org/10.1016/j.neuroimage.2015.02.050

21. Kraft, I., Schreiber, J., Cafiero, R., Metere, R., Schaadt, G., Brauer, J., … Skeide, M. A. (2016). Predicting early signs of dyslexia at a preliterate age by combining behavioral assessment with structural MRI. NeuroImage, 143, 378–386. https://doi.org/10.1016/j.neuroimage.2016.09.004

22. Langer, N., Peysakhovich, B., Zuk, J., Drottar, M., Sliva, D. D., Smith, S., … Gaab, N. (2015). White Matter Alterations in Infants at Risk for Developmental Dyslexia. Cerebral Cortex, 88C((3)), bhv281. https://doi.org/10.1093/cercor/bhv281

23. Lefly, D. L., & Pennington, B. F. (2000). Reliability and validity of the adult reading history questionnaire. Journal of Learning Disabilities, 33(3), 286–296. https://doi.org/10.1177/002221940003300306

24. Myers, C. A. C. A., Vandermosten, M., Farris, E. A. E. A., Hancock, R., Gimenez, P., Black, J. M. J. M., … Hoeft, F. (2014). White matter morphometric changes uniquely predict children’s reading acquisition. Psychological Science, 25(10), 1870–1883. https://doi.org/10.1177/0956797614544511

25. Ozernov-Palchik, O., Norton, E. S., Wang, Y., Beach, S. D., Zuk, J., Wolf, M., … Gaab, N. (2019). The relationship between socioeconomic status and white matter microstructure in pre-reading children: A longitudinal investigation. Human Brain Mapping, 40(3), 741–754. https://doi.org/10.1002/hbm.24407

26. Romeo, R. R., Segaran, J., Leonard, J. A., Robinson, S. T., West, M. R., Mackey, A. P., … Gabrieli, J. D. E. (2018). Behavioral/Cognitive Language Exposure Relates to Structural Neural Connectivity in Childhood. https://doi.org/10.1523/JNEUROSCI.0484-18.2018

27. Saygin, Z. M., Norton, E. S., Osher, D. E., Beach, S. D., Cyr, A. B., Ozernov-Palchik, O., … Gabrieli, J. D. E. (2013). Tracking the roots of reading ability: white matter volume and integrity correlate with phonological awareness in prereading and early-reading kindergarten children. The Journal of Neuroscience: The Official Journal of the Society for Neuroscience, 33(33), 13251–13258. https://doi.org/10.1523/JNEUROSCI.4383-12.2013

28. Su, M., Thiebaut de Schotten, M., Zhao, J., Song, S., Zhou, W., Gong, G., … Shu, H. (2020). Influences of the early family environment and long-term vocabulary development on the structure of white matter pathways: A longitudinal investigation. Developmental Cognitive Neuroscience, 42, 100767. https://doi.org/10.1016/j.dcn.2020.100767

29. Swagerman, S. C., van Bergen, E., Dolan, C., de Geus, E. J. C., Koenis, M. M. G., Hulshoff Pol, H. E., & Boomsma, D. I. (2017). Genetic transmission of reading ability. Brain and Language, 172. https://doi.org/10.1016/j.bandl.2015.07.008

30. Trzaskowski, M., Harlaar, N., Arden, R., Krapohl, E., Rimfeld, K., McMillan, A., … Plomin, R. (2014). Genetic influence on family socioeconomic status and children’s intelligence. Intelligence, 42(1), 83–88. https://doi.org/10.1016/j.intell.2013.11.002

31. Ursache, A., & Noble, K. G. (2016). Socioeconomic status, white matter, and executive function in children. Brain and Behavior, 6(10). https://doi.org/10.1002/brb3.531

32. van Bergen, E., van der Leij, A., de Jong, P. F., Bergen, E. van, & Leij, A. van der. (2014). The intergenerational multiple deficit model and the case of dyslexia. Frontiers in Human Neuroscience, 8, 346. https://doi.org/10.3389/fnhum.2014.00346

33. Vanderauwera, J., Wouters, J., Vandermosten, M., & Ghesquière, P. (2017). Early dynamics of white matter deficits in children developing dyslexia. Developmental Cognitive Neuroscience, 27. https://doi.org/10.1016/j.dcn.2017.08.003

34. Vanderauwera, Jolijn, De Vos, A., Forkel, S. J., Catani, M., Wouters, J., Vandermosten, M., & Ghesquière, P. (2018). Neural organization of ventral white matter tracts parallels the initial steps of reading development: A DTI tractography study. Brain and Language, 183, 32–40. https://doi.org/10.1016/j.bandl.2018.05.007

35. Vandermosten, M., Boets, B., Poelmans, H., Sunaert, S., Wouters, J., & Ghesquière, P. (2012). A tractography study in dyslexia: Neuroanatomic correlates of orthographic, phonological and speech processing. Brain, 135(3), 935–948.

36. Vandermosten, M., Cuynen, L., Vanderauwera, J., Wouters, J., & Ghesquière, P. (2017). White matter pathways mediate parental effects on children’s reading precursors. Brain and Language, 173, 10–19. https://doi.org/10.1016/j.bandl.2017.05.002

37. Vandermosten, M., Hoeft, F., & Norton, E. S. E. S. (2016). Integrating MRI brain imaging studies of pre-reading children with current theories of developmental dyslexia: A review and quantitative meta-analysis. Current Opinion in Behavioral Science, 10. https://doi.org/10.1016/j.cobeha.2016.06.007

38. Wakana, S., Caprihan, A., Panzenboeck, M. M. M., Fallon, J. J. H., Perry, M., Gollub, R. L., … Mori, S. (2007). Reproducibility of quantitative tractography methods applied to cerebral white matter. NeuroImage, 36(3), 630–644. https://doi.org/10.1016/j.neuroimage.2007.02.049

39. Wang, Y., Mauer, M. V., Raney, T., Peysakhovich, B., Becker, B. L. C., Sliva, D. D., & Gaab, N. (2017). Development of tract-specific white matter pathways during early reading development in at-risk children and typical controls. Cerebral Cortex, 27(4), 2469–2485. https://doi.org/10.1093/cercor/bhw095

40. Wang, Y., Mauer, M. V, Raney, T., Peysakhovich, B., Becker, B. L. C., Sliva, D. D., & Gaab, N. (2016). Development of Tract-Specific White Matter Pathways During Early Reading Development in At-Risk Children and Typical Controls. Cerebral Cortex (New York, N.Y.: 1991). https://doi.org/10.1093/cercor/bhw095

41. Yamagata, B., Reiss, A. L., Mimura, M., Hoeft, F., Murayama, K., Black, J. M., … Yang, T. T. (2016). Female-specific intergenerational transmission patterns of the human corticolimbic circuitry. Journal of Neuroscience, 36(4), 1254–1260. https://doi.org/10.1523/JNEUROSCI.4974-14.2016

42. Yeatman, J., & Dougherty, R. (2012). Development of white matter and reading skills. Proceedings of The. Retrieved from http://www.pnas.org/content/109/44/E3045.short

43. Yu, X., Zuk, J., & Gaab, N. (2018). What Factors Facilitate Resilience in Developmental Dyslexia? Examining Protective and Compensatory Mechanisms Across the Neurodevelopmental Trajectory. Child Development Perspectives. https://doi.org/10.1111/cdep.12293

44. Zuk, J., Dunstan, J., Norton, E., Yu, X., Ozernov-Palchik, O., Wang, Y., … Gaab, N. (2020). Multifactorial pathways facilitate resilience among kindergarteners at risk for dyslexia: A longitudinal behavioral and neuroimaging study. Developmental Science. https://doi.org/10.1111/desc.12983

