## Supplementary material for "The influence of intergenerational transfer of white matter tracts on early reading development": For results using raw scores, please see Supplementary Information

### Reading development based on raw scores of WRMT-BSC subtests

In the main text, reading development is calculated by subtracting the standardized scores of basic reading of the Woodcock Reading Mastery Tests (WRMT-BSC) in Grade 3 from the ones in Kindergarten. Since raw scores can be more intuitive to examine reading development and since WRMT_BSC is a composite score of the subtests Word Attack (WRMT-WA) and Word Identification (WRMT-WID), we provide as Supplementary Information the analyses in which reading development is calculated by subtracting the raw scores of these subtests (WRMT-WA, WRMT-WID) in Kindergarten from the ones in Grade 3. In general, the pattern of results is similar as when reading development is based on the standardized scores of WRMT-BSC.

### Reading development – white matter correlations in children

Concerning the association between *children's reading development and Arcuate Fasciculus (AF)*, substantial correlations with the right but not with the left AF were observed. More specifically, Bayesian statistics showed substantial to anecdotal evidence for no association between FA in left AF and reading development on WRMT-WID ($BF_{10}$ = 0.23) and WRMT-WA ($BF_{10}$ = 0.34). In contrast, substantial evidence was found for an association of right AF with reading development on WRMT-WID ($BF_{10}$ = 4.12) and WRMT-WA ($BF_{10}$ = 6.54). Frequentist analyses confirmed this by showing no significant correlations for left AF with reading development (WRMT-WID: $r$ = .05, $p$ = .775; WRMT-WA: $r$ = .17, $p$ = .334), while significant correlations were observed for right AF (WRMT-WID: $r$ = .43, $p$ = .013; WRMT-WA: $r$ = .46, $p$ = .008).

Concerning the association between *children's reading development and Inferior Fronto-Occipital-Fasciculus (IFOF)*, correlations were found for bilateral IFOF but were stronger in right IFOF. More specifically, there was anecdotal to substantial evidence for an association between FA in left IFOF and reading development on WRMT-WID ($BF_{10}$ = 3.47) and WRMT-WA ($BF_{10}$ = 1.58). For right IFOF, substantial to very strong evidence was found for an association between right IFOF and reading development on WRMT-WID ($BF_{10}$ = 41.68) and WRMT-WA ($BF_{10}$ = 5.65). Frequentist analyses displayed significant correlations between reading development and left IFOF (WRMT-WID: $r$ = .41, $p$ = .016; WRMT-WA: $r$ = .36, $p$ = .042), as well as significant correlations with right IFOF (WRMT-WID: $r$ = .55, $p$ < .001; WRMT-WA: $r$ = .45, $p$ = .009).

In summary, Bayesian and frequentist analyses showed clear correlations (defined as $BF_{10} > 3$ and $p < .05$) for children's reading development and FA in right AF and right IFOF (see Supplementary Figure 1), although some evidence was also found for correlations between reading development and left IFOF.

*Supplementary figure 1: Correlations between children's reading development and FA in white matter tracts. Only correlations for which Bayesian statistics showed at least substantial evidence (BF > 3) and frequentist statistics showed significant results (p < .05) are depicted.*

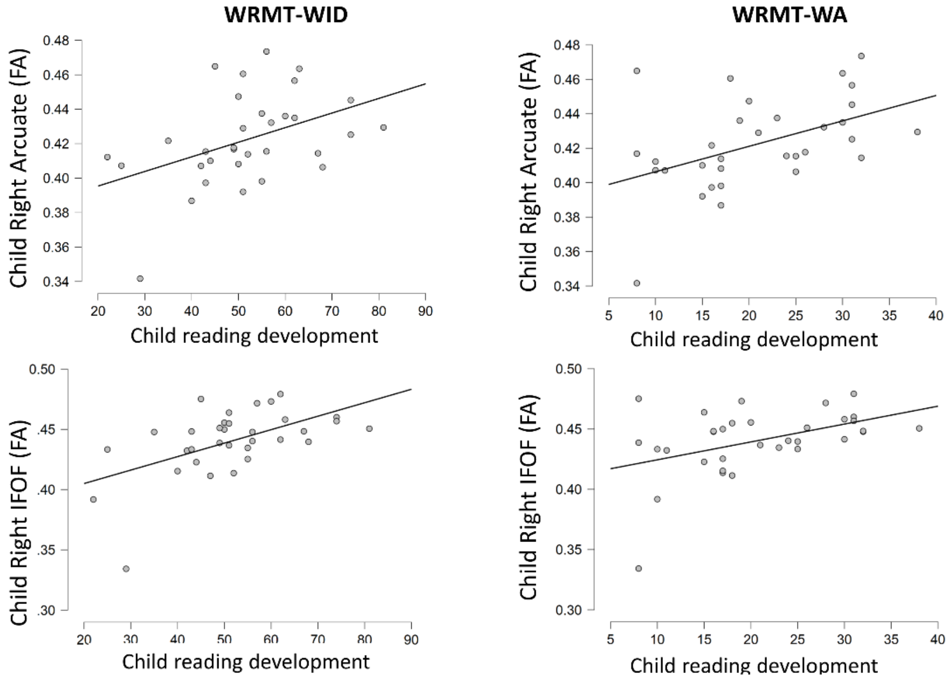


### Mediation Analyses

Mediation analyses showed that the effect of children's *right IFOF* on reading development was largely explained by mother's and father's right IFOF. More specifically, for *WRMT-WID*, the total effect, i.e., the regression coefficient between child's right IFOF and reading development, was 20.15 and significant ($z$ = 3.78, $p$ < .001), which is in line with the correlational analyses reported above. However, when the mediation effect (i.e. parental FA in right IFOF) was taken into account, the direct effect of child's right IFOF on reading development decreased to 4.31 and became non-significant ($z$ = 0.37, $p$ = .715). The indirect effect was (19)*(.50) = 9.50 for mothers and (26)*(.24) = 6.35 for fathers. This implies that the indirect effects together (15.85) explained 79% of the total effect of child's right IFOF on reading development. However, when we tested the significance of these indirect effects using bootstrapping procedures (unstandardized indirect effects were computed for each 1000 bootstrapped sample), the total indirect effects did not reach significance ($p$ = .155, 95% confidence interval range: -7.36 to 44.27). For *WRMT-WA*, a similar pattern was observed. The total effect of child's right IFOF on reading development was 16.39 and significant ($z$ = 2.87, $p$ = .004). However, when parental FA in right IFOF was taken into account as a mediator, the direct effect of child's right IFOF on reading development decreased to 7.87 and became non-significant ($z$ = 0.62, $p$ = .534). The indirect effect was (18)*(.0.18) = 3.21 for mothers and (26)*(.20) = 5.31 for fathers. This implies that the indirect effects together (8.52) explained 52% of the total effect of child's right IFOF on reading development. However, using bootstrapping procedures the total indirect effect was again not significant (*p* = .457, 95% confidence interval range: -22.36 to 56.67).

Concerning *left IFOF*, mediation analyses showed that the effect of child's left IFOF on reading development was largely explained by mother's and father's left IFOF. More specifically, for *WRMT-WID*, the total effect of child's left IFOF and reading development was 14.54 and significant ($z$ = 2.63, $p$ = .009). However, when the mediation effect (i.e. parental FA in left IFOF) was taken into account, the direct effect of child's left IFOF on reading development decreased to 4.52 and became non-significant ($z$ = 0.45, $p$ = .653). The indirect effect was (13)*(.35) = 4.62 for mothers and (30)*(.18) = 5.40 for fathers. This implies that the indirect effects together (10.02) explained 69% of the total effect of child's left IFOF on reading development. However, total indirect effects using bootstrapping procedures did not reach significance ($p$ = .268, 95% confidence interval range: -8.47 to 28.39). For *WRMT-WA*, the total effect of child's left IFOF on reading development was 12.47 and significant ($z$ = 2.19, $p$ = .028). Again, when parental FA in left IFOF was taken into account as a mediator, the direct effect of child's left IFOF on reading development decreased to 4.50 and became non-significant ($z$ = 0.39, $p$ = .697). The indirect effect was (14)*(.11) = 1.52 for mothers and (30)*(.22) = 6.45 for fathers. This implies that the indirect effects together (7.97) explained 64% of the total effect of child's left IFOF on reading development. However, using bootstrapping procedures the total indirect effect was not significant ($p$ = .437, 95% confidence interval range: -13.77 to 30.70).
